# Robust collection and processing for label-free single voxel proteomics

**DOI:** 10.1101/2023.08.14.553333

**Authors:** Reta Birhanu Kitata, Marija Velickovic, Zhangyang Xu, Rui Zhao, David Scholten, Rosalie K. Chu, Daniel J. Orton, William B. Chrisler, Jeremy V. Mathews, Paul D. Piehowski, Tao Liu, Richard D. Smith, Huiping Liu, Clive H. Wasserfall, Chia-Feng Tsai, Tujin Shi

## Abstract

With advanced mass spectrometry (MS)-based proteomics, genome-scale proteome coverage can be achieved from bulk tissues. However, such bulk measurement lacks spatial resolution and obscures important tissue heterogeneity, which make it impossible for proteome mapping of tissue microenvironment. Here we report an integrated wet collection of single tissue voxel and Surfactant-assisted One-Pot voxel processing method termed wcSOP for robust label-free single voxel proteomics. wcSOP capitalizes on buffer droplet-assisted wet collection of single tissue voxel dissected by LCM into the PCR tube cap and MS-compatible surfactant-assisted one-pot voxel processing in the collection cap. This convenient method allows reproducible label-free quantification of ∼900 and ∼4,600 proteins for single voxel from fresh frozen human spleen tissue at 20 μm × 20 μm × 10 μm (close to single cells) and 200 μm × 200 μm × 10 μm (∼100 cells), respectively. 100s-1000s of protein signatures with differential expression levels were identified to be spatially resolved between spleen red and white pulp regions depending on the voxel size. Region-specific signaling pathways were enriched from single voxel proteomics data. Antibody-based CODEX imaging was used to validate label-free MS quantitation for single voxel analysis. The wcSOP-MS method paves the way for routine robust single voxel proteomics and spatial proteomics.

## INTRODUCTION

The human body comprises trillions of cells with hundreds of different cell types, which are spatially organized along with their extracellular matrix to form tissues and organs with diverse functionalities^1, 2^. A tissue is an ensemble of certain types of cells spatially located at distinct anatomical regions to execute region-specific functions^3^. There is significant heterogeneity across different anatomical regions within an organ tissue^4-6^. Therefore, spatial proteomics for enabling precise, comprehensive, proteomic mapping of human tissues at high spatial resolution is crucial for phenotypic characterization of tissue heterogeneity and microenvironment within spatial context to improve our understandings of tissue biology and how unique cell types contribute to heterogeneous tissues^7^.

Antibody-based immunoassays such as immunohistochemistry (IHC)^8, 9^, immunofluorescence (IF)^9^, CODEX (codetection by indexing)^10, 11^, MIBI (multiplexed ion beam imaging)^12, 13^, and mass cytometry^14^ are primarily used for single-cell or near single-cell resolution spatial targeted proteomics mapping of human tissues^15^. However, they require a prior knowledge with the limited ability for discovery and only can open a small window into the tissue microenvironment due to their inherent limitations (e.g., low multiplex with up to 60 protein markers, and long lead time and difficulties in generating high-quality antibodies for spatial proteomics imaging). Unlike antibody-based targeted proteomics, mass spectrometry (MS)-based global proteomics enables for reliable simultaneous detection and quantification of >10,000 proteins from bulk tissue samples with a large number of mixed populations of cells (e.g., ≥10 million of cells ≈ 1 mg of tissues)^16-19^. Such bulk measurements average out cell-type specific tissue heterogeneity when cells of interest only account for a small portion of the total cell populations. Thus, they cannot provide any insights into tissue heterogeneity and microenvironment within spatial context for understanding functional spatial biology.

Laser capture microdissection (LCM) is routinely used for microscopic isolation and capture of subpopulations of cells, individual cells, or regions of interest in a complex heterogenous tissue section^20-25^. In past two decades a surge of papers was reported using LCM for tissue spatial molecular profiling^24^. When LCM collection was combined with advanced MS-based proteomics, LCM-MS allows for region- and cell type-resolved proteome characterization of human tissues^23, 25^. Currently, the combined LCM and cap-assisted methods are most widely used for convenient spatial proteomics due to their cost-effective, easy implementation and an easy-to-use tissue voxel collection and processing workflow^21, 23, 26^. They collect LCM-dissected tissue voxels on the cap, transfer the collected voxels or tissue voxel lysates to the tube, and process them in the tube. For example, Davis *et al*. performed a series of voxel sample preparation optimizations and developed a sensitive cap-assisted SP3 method for enabling identification of ∼1,500 proteins from as little as 100 Purkinje cells dissected by LCM with the total area of 0.06 mm^2^ at a thickness of 10 μm^27^. Drummond *et al*. developed a wet cap-assisted localized proteomics method for quantitative cell type-specific proteomics in Alzheimer’s disease (AD) research^28, 29^. 1000s of neurons, amyloid plaques or neurofibrillary tangles (NFTs) (the total area of 0.5-2.5 mm^2^) from AD patient brain tissues dissected by LCM were collected to the tube cap prefilled with distilled water and then transferred to the tube by centrifugation for sample processing^28, 30-32^. ∼500 and ∼900 proteins were identified from dissected neurons (NFTs) and amyloid plaques, respectively. By comparative analysis of neighboring non-diseased tissue voxels, this localized proteomics method allows pathway and network enrichment analysis for discovery of potential therapeutic targets and biomarkers for AD. However, current cap-assisted methods are only suitable for collection and processing of 100s-1000s of pooled tissue voxels or large regions of interest. They cannot be used for effective single voxel proteomics due to significant sample loss during transferring and processing of single tissue voxel at small sizes (e.g., 0.0004-0.04 mm^2^ equivalent to the area of 1-100 cells). The ability for robust proteomic analysis of single voxel (pixel) (i.e., single voxel (pixel) proteomics) is essential for high spatial resolution proteome mapping of human tissues and comprehensive characterization of tissue microenvironment. Furthermore, single voxel proteomics is crucial for analysis of few cells of interest within the complex tissue microenvironment (e.g., few tumor cells in biopsy tissue at the early cancer stage) with great potential of moving towards precision medicine.

To alleviate the shortcomings of existing spatial proteomics methods, we have recently developed a broadly adoptable robust cap-assisted method for quantitative label-free single voxel proteomics. This method capitalizes on both wet collection and Surfactant-assisted One-Pot processing of single tissue voxel termed wcSOP on the tube cap prefilled with a cocktail of buffer containing DDM surfactant and protease (**Fig. 1a**). wcSOP greatly reduces sample transfer and surface adsorption losses, thus significantly improving detection sensitivity for MS analysis of single tissue voxel at a small size. wcSOP-MS was demonstrated to enable reliable label-free quantification of ∼900 (∼4,600) proteins from single tissue voxel at 0.0004 mm^2^ (0.04 mm^2^) at a thickness of 10 μm with standard LC-MS platforms. wcSOP-MS enabled to identify spatially resolved proteome changes and enriched pathways at high spatial resolutions. These results demonstrate great potential of wcSOP-MS for broad applications in the biomedical research. A commonly and easily accessible Q Exactive Plus or HF MS platform was used for the development of wcSOP-MS and its application demonstration.

**Fig. 1.**
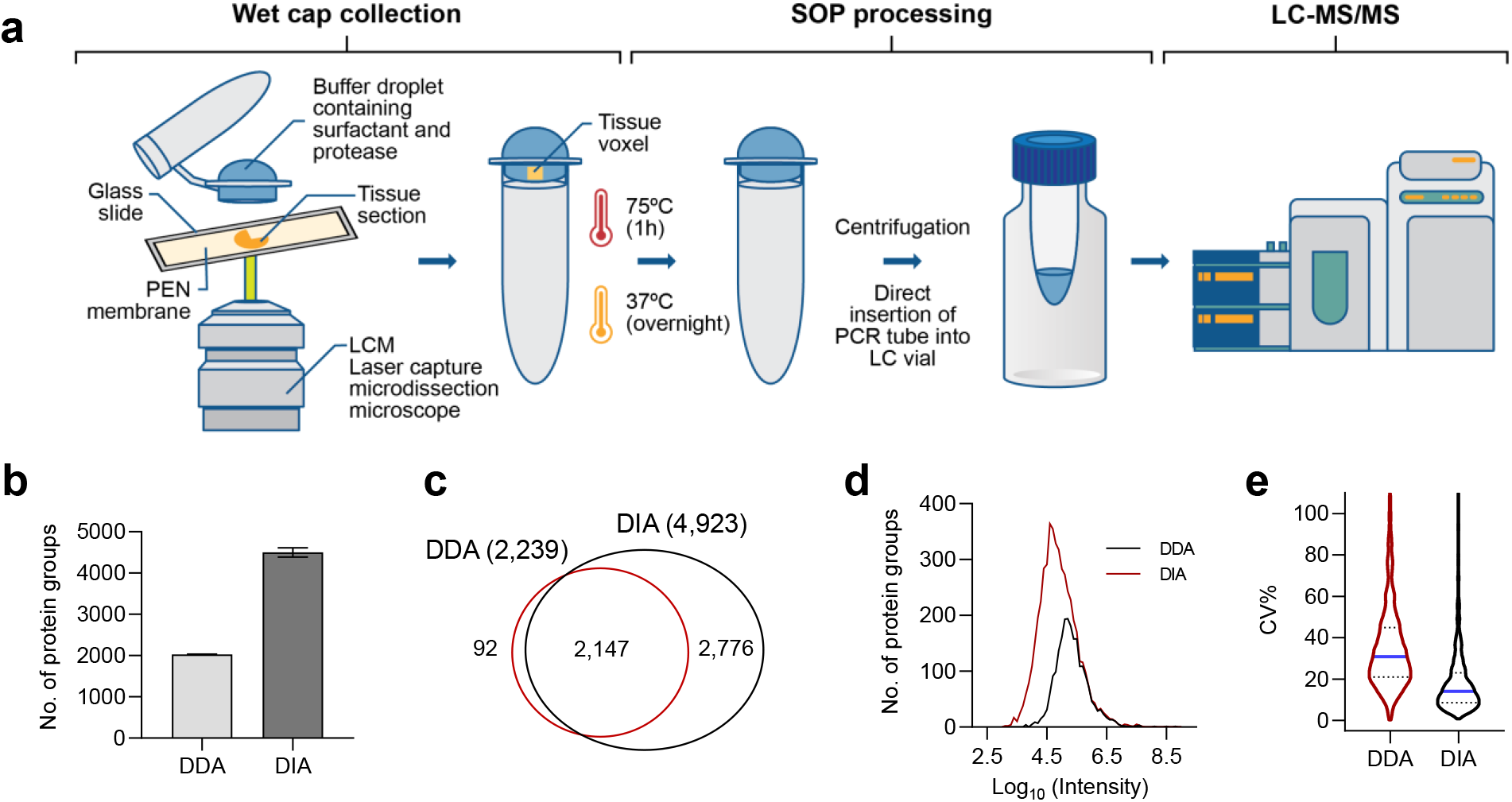
Schematic diagram of the wcSOP-MS workflow. **a**, There are three steps for wcSOP-MS: wet cap collection, SOP processing, and LC-MS/MS analysis. A tissue slide mounted on PEN membrane is cut by laser and single tissue voxel is catapulted to a dome-shaped PCR tube cap prefilled with a cocktail of buffer containing DDM surfactant for tissue lysis and a mixture of Lys-C and trypsin protease for protein digestion. Single voxel is incubated at 75ºC for 1 h (tissue lysis and protein denaturation) and 37ºC for overnight (protein digestion) on the cap at a hanging droplet position to have full interaction between the voxel and the cocktail buffer. After voxel processing, the digested peptides are transferred to the bottom of the tube by centrifugation at 2000 *g* for 5 min. Prior to LC-MS analysis, the cap of the PCR tube is removed and the tube is inserted into a sample vial to avoid sample transfer loss. Single tissue voxel is analyzed by using standard LC-MS platform for quantitative proteomic analysis. The freely-available open-source MaxQuant and DIA-NN software tools are used for DDA and DIA label-free quantitation, respectively. **b**, Comparison of the number of identified protein groups (No. of protein groups) between DDA and DIA for single spleen tissue voxel at a size of 200 μm × 200 μm × 10 μm. **c**, Venn diagram of protein overlap between DDA and DIA for single spleen tissue voxel at a size of 200 μm × 200 μm × 10 μm. **d**, Proteome dynamic range for DDA and DIA quantification of single spleen tissue voxel at a size of 200 μm × 200 μm × 10 μm. **e**, Comparison of the coefficient of variations (CVs) between DDA and DIA for analysis of single spleen tissue voxel at a size of 200 μm × 200 μm × 10 μm.

## RESULTS

### Systematic evaluation of cap-assisted methods for tissue voxel collection and processing

The combined LCM and various cap-assisted methods are most widely used for convenient spatial proteome profiling of subpopulations of cells, image-guided regions of interest, and tissue microenvironment from heterogenous tissues due to their cost-effective, easy implementation without the need of any special devices or a highly skilled person^21, 23, 26, 33-35^. Commercially available adhesive tube caps or standard PCR tube caps are used for collection and processing of tissue voxels dissected by LCM. However, current cap-assisted methods are only suitable for collection and processing of 100s-1000s of pooled tissue voxels with the total area of ∼0.5-2.5 mm^2^ at relatively large processing volumes. They cannot be used for high-resolution single voxel proteomics because there is a technical challenge in effective transferring and processing of single tissue voxel at small sizes (e.g., 0.0004-0.04 mm^2^ equivalent to the area of 1-100 cells). Single voxel proteomics is crucial for precise spatial characterization of tissue heterogeneity and generation of high spatial resolution tissue maps.

To address this issue, we have recently systematically evaluated and optimized cap-assisted methods for single voxel proteomics. Individual spleen tissue voxels at the size of 0.04 mm^2^ (200 μm × 200 μm) with a thickness of 10 μm were collected into the tube caps, processed by our previously developed SOP, and analyzed by data dependent acquisition (DDA) MS analysis. There are two types of cap-assisted collection methods: dry collection with adhesive polymer attached to the cap and wet collection with prefilled water or buffer on the cap. We first evaluated a conventional dry cap-assisted collection and processing method for single voxel proteomics. Tissue lysis occurs on the cap, and protein denaturation and digestion are performed in the tube. Different lysis buffer volumes on the cap were evaluated, and high irreproducibility was observed with protein identification from 715 to 1,432. This irreproducibility is most likely due to incomplete tissue lysis and/or significant adsorption loss to adhesive polymer. We next evaluated all tissue lysis, denaturation, and digestion on the cap (i.e., ‘all-in-one’ dry cap-assisted SOP termed dcSOP) with the processing volume of 10 μL (the maximal volume for the adhesive cap). The selection of 10 μL processing volume is to fully cover single tissue voxel since it may be collected at the edge of cap. The technical reproducibility was greatly improved with 1,724 proteins and CV<10%. However, there are two drawbacks for dcSOP, adhesive polymer detached from the cap during centrifugation and difficulty in tightly sealing the adhesive tube cap. They can severely affect its robustness in sample processing and injection for single voxel proteomics analysis.

In parallel, we have revisited a conventional wet cap-assisted method used for spatial proteomics. Prior to voxel collection, 25 μL buffer was added to the dome-shaped PCR tube cap with the maximal volume of ∼25 μL to form the droplet with the largest surface area at a diameter of ∼5.4 mm (**Fig. 1a**). The large droplet ensures precise convenient collection of single tissue voxel with ∼100% success rate. After collection the PCR tube was centrifuged at 2000 *g* to presumably transfer the tissue voxel from the cap to the bottom of the tube. The collected single voxel was directly processed in the tube using SOP. To maximize sample recovery and processing reproducibility, we have optimized a series of SOP conditions including different types of buffers, different buffer volumes, and different temperatures for tissue lysis and protein denaturation. Unfortunately, low reproducibility was consistently observed no matter what conditions were used. With demonstrated performance of SOP-MS for robust single-cell proteomics analysis, the only explanation is that there is an issue for successfully transferring of the collected voxel from the cap to the bottom of the well. This was confirmed by close voxel tracing during the transferring step. Microscope was used to monitor the entire voxel transfer process (voxel collection, cap checking after centrifugation, and voxel checking in PCR tube after drying in a SpeedVac concentrator). The collected single voxel was found almost everywhere such as on the cap, on the top, the middle, the low side tube wall, and at the bottom of the tube. The probability of single voxel positions is widely distributed though there is a much higher chance for the collected single voxel on the bottom of the tube, which results in low processing reproducibility for single voxel proteomics. This can be attributed to much stronger electrostatic interaction between the small tissue voxel and the plastic tube wall or cap than the ultracentrifugation force for single small tissue voxel.

With an insurmountable technical challenge in transferring single voxel down to the well bottom as well as our experience in dcSOP we turned to pursue a different way for the wet cap-assisted SOP method for both collection and processing of single voxel on the cap. The wet cap-assisted SOP method termed wcSOP capitalizes super tight sealing of the dome-shaped PCR tube cap, low-bind PCR tube cap, and precise collection of single tissue. We evaluated three different types of wcSOP methods (i.e., 3-step, 2-step, and ‘all-in-one’ wcSOP) based on the number of steps for the addition of buffers to the cap for voxel collection, tissue lysis, and protein digestion. After the buffer addition, PCR tube was in the normal position with the collected voxel hanging on the cap. With the use of 25 μL buffer for collection of single voxel, the total processing volume is ∼22 μL due to partial water evaporation to the tube wall. The voxel processing volume is in the range of 5-50 μL previously used for SOP processing of 1-100 human cells^36, 37^. All the three wcSOP methods enabled for identification of >10,000 unique peptides and >2,100 protein groups with high reproducibility from single human spleen voxel at the size of 0.04 mm^2^ (200 μm × 200 μm). As the buffer addition steps reduced the number of identified protein groups (unique peptides) were increased with ∼2,200 protein groups (∼11,100 unique peptides) for ‘all-in-one’ wcSOP. Consequently, the total area for the extracted ion chromatogram (XIC) was also increased from 3-step to ‘all-in-one’ wcSOP methods. Much less variation in the XIC area was observed for ‘all-in-one’ wcSOP, suggesting higher processing reproducibility. Venn diagram plotting has shown high overlap of protein groups (∼88%) identified from the three wcSOP methods. The number of identified protein groups (unique peptides) from wcSOP is much higher than that from dcSOP presumably due to more effective contact surface interaction between the collected voxel and buffers during sample processing for increased sample recovery. With high sample recovery from all the evaluation results as well as easy convenient operation, ‘all-in-one’ wcSOP was selected as the optimal cap-assisted voxel collection and processing for single voxel proteomics analysis.

### ‘All-in-one’ wcSOP-MS for sensitive label-free single voxel proteomics

With systematic evaluation of different cap-assisted voxel collection and processing, the ‘all-in-one’ wcSOP-MS method was selected for robust single voxel proteomics analysis (**Fig. 1a**). In the ‘all-in-one’ wcSOP-MS workflow, a cocktail of buffer containing DDM surfactant for tissue lysis and protease (the mixture of lysin C and trypsin) for protein digestion is added to the dome-shaped PCR tube cap to form a large droplet with a diameter of ∼5.4 mm, which warrants robust precise collection of single tissue voxel. The collected tissue voxel hangs in the cap to maximize the contact surface interaction between the voxel and the cocktail buffer for effective tissue lysis and digestion (**Fig. 1a**). Then it is processed on the cap without sample transfer using our recently developed SOP method for single cell proteomics. After processing the tissue voxel digest is transferred to the bottom of PCR tube by centrifugation followed by direct LC-MS analysis.

To improve proteome coverage for single voxel proteomics we have systematically compared label free data-independent acquisition (DIA) with DDA for analysis of different sizes of tissue voxels derived from fresh frozen human spleen tissue. With recent advances in MS instrumentation and informatics tools for DIA data analysis, label free DIA MS analysis has gained popularity for rapid, reproducible, in-depth proteome profiling. Tissue voxels at the sizes of 0.0004 mm^2^ (20 μm × 20 μm), 0.0025 mm^2^ (50 μm × 50 μm), 0.01 mm^2^ (100 μm × 100 μm), and 0.04 mm^2^ (200 μm × 200 μm), which are equivalent to single cells, 6 cells, 25 cells, and 100 cells, respectively, were collected and processed using wcSOP followed by DIA or DDA MS analysis with 4 biological replicates per condition. For DDA MS analysis, 344 ± 99, 622 ± 172, 1542 ± 39 and 2051 ± 18 protein groups were identified from single voxel at the sizes of 0.0004 mm^2^, 0.025 mm^2^, 0.01 mm^2^ and 0.04 mm^2^, respectively, while DIA MS analysis allows for identification of 871 ± 51, 1948 ± 228, 4052 ± 111, 4500 ± 8 protein groups (**Fig. 1b**). Such voxel analysis represents one of the deepest proteome coverages that can be achieved for small tissue voxels. While DIA covers almost all of proteins identified by DDA, protein abundance distribution shows DDA detected only those in high abundant region with lower quantitation reproducibility (**Figs. 1c-1d**). This suggests that DIA MS analysis can provide much higher proteome coverage with ∼2.5-3-fold more protein groups than DDA MS analysis. High reproducibility with lower CV% was observed for DIA MS analysis (**Fig. 1e**). Unless otherwise noted, DIA MS analysis was used for later voxel data generation to evaluate wcSOP-MS performance on different types of tissues.

### Evaluation of wcSOP-MS performance using human spleen tissue

Fresh (snap) frozen tissue is often the first choice for MS-based proteomics analysis as well as for evaluation of method or platform performance because of its higher protein yield and less sample processing steps than other types of tissues. Different sizes of tissue voxels from fresh frozen spleen tissue were selected to evaluate wcSOP-MS performance. The spleen is the largest secondary lymphoid organ which contains highly organized and mainly compartmentalized structure of the red pulp (RP) through which the blood is continuously filtered to remove old or damaged red blood cells and the white pulp (WP) responsible for immune surveillance against pathogens^38^. The spleen immune system plays a significant role in innate and adaptive immunity^39^. Thus, the two distinct regions (RP and WP) known to have different cell types are an ideal model to assess wcSOP-MS performance on fresh frozen tissues.

Spleen tissue voxels at the sizes of 0.0004 mm^2^, 0.0025 mm^2^, 0.01 mm^2^, and 0.04 mm^2^ from both RP and WP regions were collected and processed by wcSOP followed by DIA MS analysis. wcSOP-MS allows for identification of 709 ± 300, 2252 ± 42, 3630 ± 208 and 4673 ±40 protein groups from single WP voxels at the sizes of 0.0004 mm^2^, 0.025 mm^2^, 0.01 mm^2^ and 0.04 mm^2^, respectively (**Fig. 2a**). Similarly, 881 ± 51, 1959 ± 227, 4061 ± 385, 4509 ± 40 protein groups can be identified from single RP voxels at the sizes of 0.0004 mm^2^, 0.025 mm^2^, 0.01 mm^2^ and 0.04 mm^2^, respectively (**Fig. 2a**). >3,000 and >30,000 unique peptides were identified from voxels at the sizes of 0.0004 mm^2^ and 0.04 mm^2^, respectively (**Fig. 2a**). This strongly suggests high reproducibility of wcSOP-MS for label free proteome profiling of small tissue voxels.

**Fig. 2.**
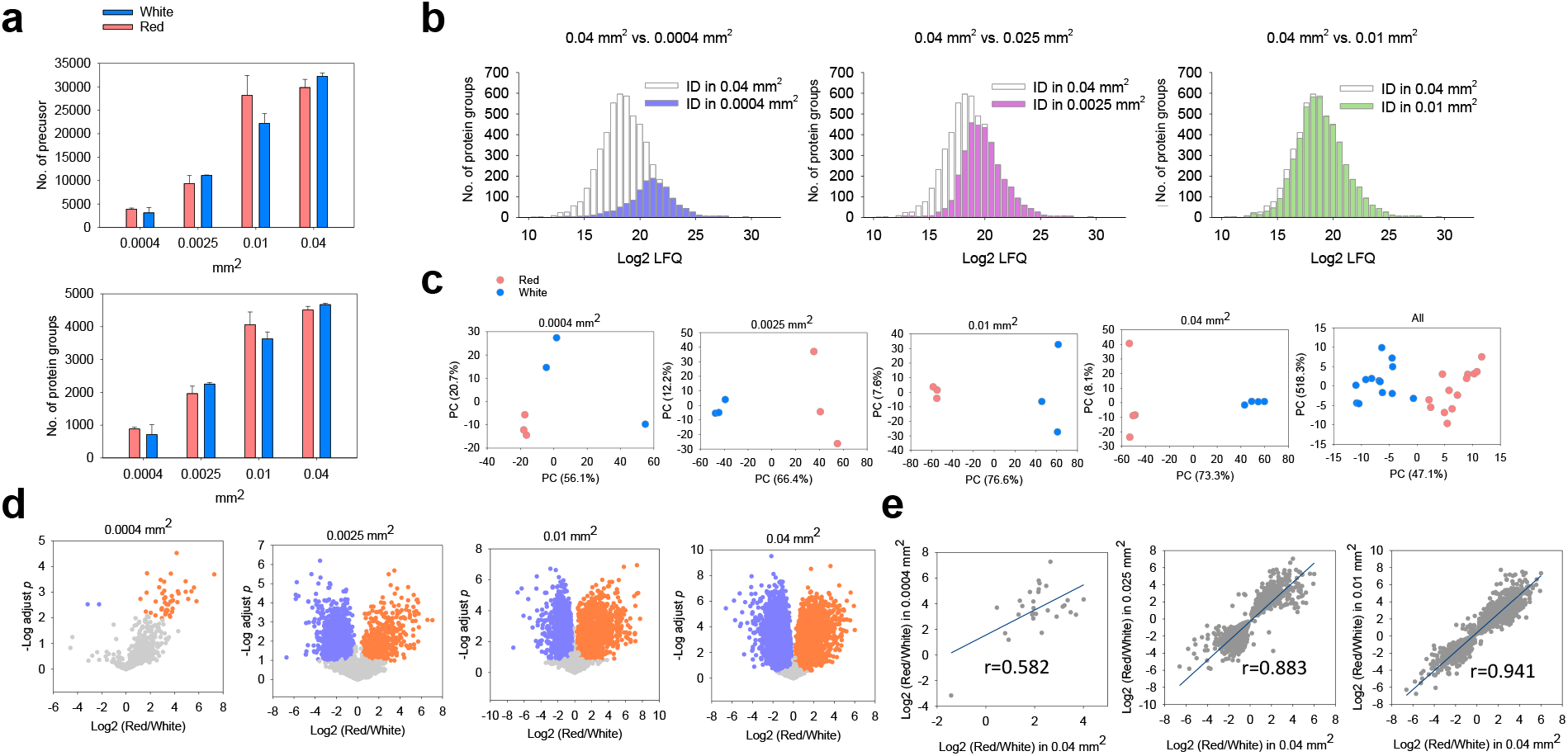
wcSOP-MS analysis of different sizes of single voxel from red and white pulp regions of human spleen tissue. **a**, The number of identified unique peptides and protein groups at different sizes of single voxel. **b**, Proteome dynamic range from DIA quantification of single voxel at different sizes against the largest voxel size of 0.04 mm^2^. **c**, PCA analysis of the commonly expressed protein abundance between single voxels from the red and white pulp regions at different sizes. **d**, Volcano plots between single voxels from the red and white pulp regions at different sizes. **e**, Correlation analysis of the protein abundance ratio of the red over white pulp regions from small single voxels with the largest single voxels at 0.04 mm^2^. Single tissue voxels at 0.0004 mm^2^ (20 μm × 20 μm equivalent to single cells), 0.0025 mm^2^ (50 μm × 50 μm equivalent to 6 cells), 0.01 mm^2^ (100 μm × 100 μm equivalent to 25 cells) and 0.04 mm^2^ (200 μm × 200 μm equivalent to 100 cells) with a thickness of 10 μm were collected, processed, and analyzed using wcSOP-DIA MS with three biological replicates per condition.

For the largest voxels at 0.04 mm^2^ with the deepest proteome coverage, ∼6 orders of magnitude dynamic concentration range was observed, reflecting high detection sensitivity of wcSOP-MS even though a commonly accessible MS platform was used. As expected, most proteins identified from voxels at 0.0004 mm^2^ are highly abundant (**Fig. 2b**). As the voxel size increased to 0.025 mm^2^, moderate abundance proteins were detected, and for voxels at 0.01 mm^2^ low abundance proteins can be identified (**Fig. 2b**). Principal component analysis (PCA) of the WP and RP regions has shown distinct separation of the two regions based on protein expression alone even for ultrasmall voxels at 0.0025mm^2^ (**Fig. 2c**). With an increase of voxel size the separation efficiency was gradually increased primarily due to the increased number of differentially expressed proteins (i.e., signatures or biomarkers) (**Fig. 2c**). Volcano plots have shown 47, 1189, 2499, and 3814 protein markers with significantly differential expression between the WP and RP regions for 0.0004 mm^2^, 0.0025 mm^2^, 0.01 mm^2^, and 0.04 mm^2^, respectively (**Fig. 2d**). To evaluate quantitation accuracy, we performed Pearson correlation analysis by comparing the ratio of the RP over WP protein abundance at smaller voxel sizes with the largest voxel size of 0.04 mm^2^. High Pearson correlation coefficients of 0.88 and 0.94 were observed for voxel sizes at 0.0025 mm^2^ and 0.01 mm^2^, respectively. But for the smallest voxels at the size of 0.0004 mm^2^ there is positive correlation but with a moderate correlation coefficient (**Fig. 2e**). This can be attributed to unreliable protein quantitation due to the lack of sufficient sensitivity for ultrasmall tissue voxels. In addition, we have compared our spleen tissue voxel data with one recent bulk study for in-depth proteome map of the human body including spleen tissue. ∼90% of proteins from spleen tissue voxels were observed in the spleen proteome with the total ∼1,1000 proteins^40^. Among these overlapped proteins 274 proteins are spleen tissue specific proteins, which accounts for 81% of all spleen tissue specific proteins^40^. All these results demonstrate the reproducibility, quantitation dynamic range, and quantitation accuracy of wcSOP-MS for robust single voxel proteomics analysis.

### Single voxel proteomics for revealing region-specific markers and signaling pathways

Using wcSOP-DIA for single voxel proteomics analysis, 1,189, 2,499, and 3,814 of protein signatures (biomarkers) can be identified for enabling differentiation of two distinct red and white pulp regions at the tissue voxel sizes of 0.025 mm^2^, 0.01 mm^2^, and 0.04 mm^2^, respectively (**Fig. 2d**). They include many cell surface CD protein markers (**Fig. 3a**), and some CD markers are known to be specific to certain cell types within the two regions (e.g., B cell markers: CD20, CD22; T cell markers: CD4, CD8A; macrophage marker: CD163). CD20, CD22 and CD44 were upregulated in the white pulp region, whereas CD8A, CD4, CD47 and CD163 upregulated in the red pulp region (**Fig. 3a**). This observation is consistent with region-specific cell type distribution and function difference between the red and white pulp regions.

**Fig. 3.**
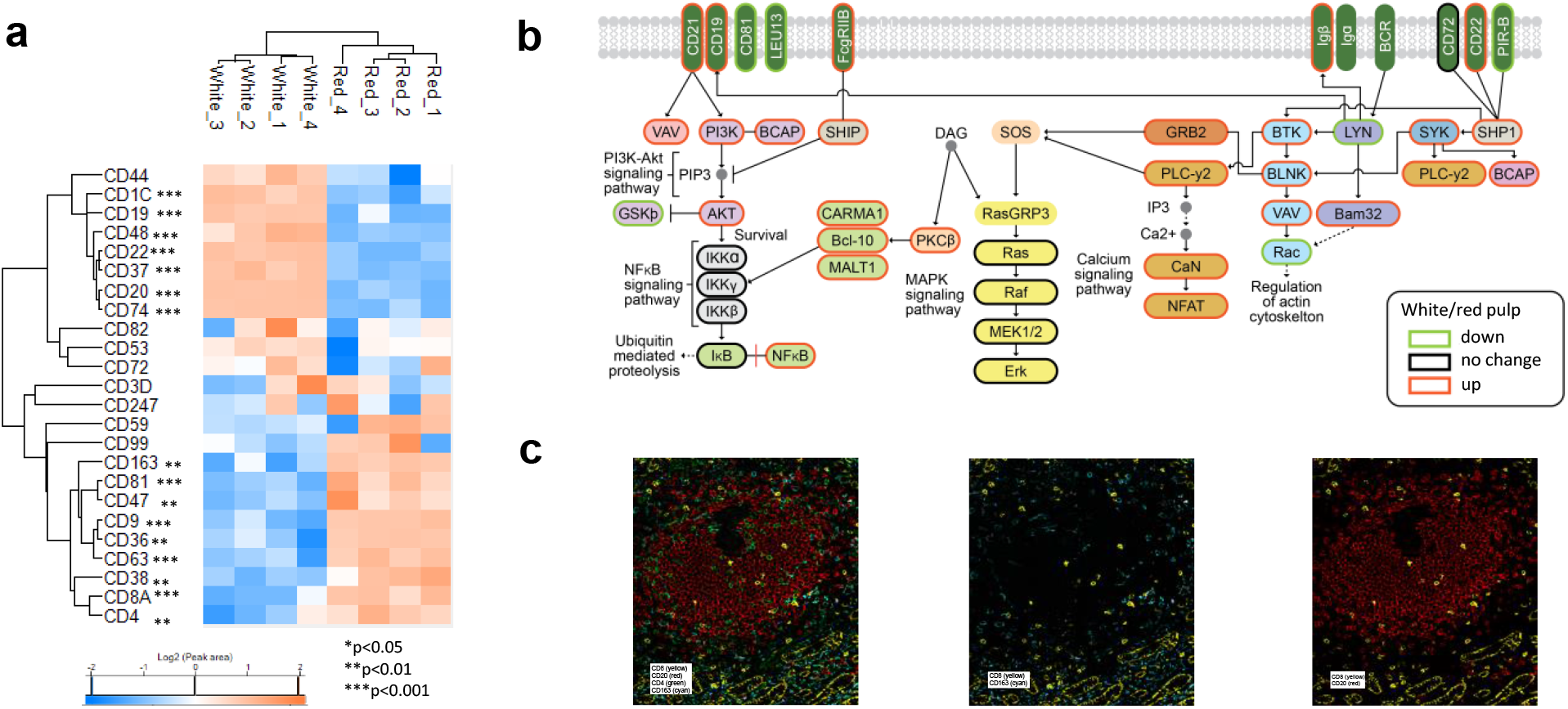
Functional analysis of single voxel proteomics data from the largest voxels. **a**, Differentially expressed CD protein markers between the red and white pulp regions. **b**, Enriched signaling pathways between the red and white pulp regions. **c**, Antibody-based CODEX image of human spleen tissue slide to validate label-free DIA MS quantitation: CD8 (yellow), CD20 (red), CD4 (green), and CD163 (cyan).

Red pulp is a loose spongy tissue with chords of reticular cells located between venous sinuses that primarily contain erythrocytes, lymphocytes, macrophages, granulocytes, and plasma cells. Unique to human spleen red pulp, CD8A lines these sinuses, termed littoral cells^41^. Senescent erythrocytes are destroyed by macrophages with hemoglobin:haptoglobin complexes being loaded onto CD163 positive macrophages for metabolic processing. Thus, CD8A, haptoglobin and CD163 were observed to have higher expression levels in the red pulp regions (**Fig. 3a**). By contrast, as a B cell specific membrane protein and an attractive target for therapeutic antibodies^42^, CD20 protein has higher expression level in the white pulp region. All these results demonstrate that single voxel proteomics enables to precisely reveal functionally important proteins across different regions at high spatial resolutions.

We have further performed KEGG pathway analysis of differentially expressed proteins between the red and white pulp regions. Several signaling pathways were enriched and they are closely related to the white or red pulp functions. One example is the B-cell receptor (BCR) signaling pathway, which is known to be crucial for normal B cell development and adaptive immunity as well as for B-cell malignancies^43^. B cell activation begins with initiation of signaling pathways (e.g., NFAT, NF-κB, and MAPK) and endocytosis of the BCR-antigen complex^44^. Our single voxel proteomics analysis can cover 56 BCR pathway proteins among the total 78 proteins with 30 proteins upregulated and 8 proteins downregulated in the white pulp region (**Fig. 3b**). One important upregulated protein in the BCR pathway is spleen tyrosine kinase (SYK), which is known to have a crucial role in adaptive immune receptor signaling^45^. SYK was a cytosolic non-receptor protein tyrosine kinase recognized as a critical element in the BCR signaling pathway and its phosphorylation activates downstream signaling pathways. Several oral Syk inhibitors are being assessed in clinical trials for multiple malignancies^46^. This suggests that with its deep proteome coverage wcSOP-based single voxel proteomics has the potential for high spatial resolution mapping of signaling pathways in human tissues to improve our understanding of tissue microenvironment.

### Validation of label-free MS quantification using CODEX imaging

As mentioned above many cell surface CD protein markers can be detected and quantified from human spleen tissue voxels using wcSOP-MS. To further validate label-free DIA MS quantitation, we have performed an antibody-based multiplexed CODetection by indexing (CODEX) approach for imaging four representative region-specific protein markers, CD20, CD4, CD8A, and CD163. CODEX imaging of human spleen tissue has shown that CD20 is highly expressed in the white pulp region, whereas CD4, CD8A, and CD16 have higher expression levels in the red pulp region (**Fig. 3c**). The results from CODEX imaging are consistent with the data from label-free DIA MS quantification (**Figs. 2d** and **3a**), which confirmed reliable single voxel proteomics analysis with wcSOP-MS.

## CONCLUSION

In summary, we report an easily implementable wcSOP-MS method that capitalizes on wet cap collection of single tissue voxel and one-pot voxel processing for robust, sensitive, label-free single voxel proteomics. With its convenient features (e.g., low-cost PCR tube cap and surfactant) as well as its being easy to use, wcSOP-MS can be readily implemented in any MS laboratory for spatial proteomics when LCM and MS are available. Its quantitation accuracy was validated with traditional antibody-based imaging methods. wcSOP-MS opens an avenue for routine single voxel proteomics and spatial proteomics.

## Acknowledgments

This work was supported by a UH3CA256967 grant (to T.S.) from the National Institutes of Health (NIH) Common Fund, Human Biomolecular Atlas Program (HuBMAP) grant, NIH RF1MH128885 (to T.S.), and NIH R01GM139858 (to T.S.). PNNL is a multi-program national laboratory operated by Battelle for the Department of Energy (DOE) under Contract DE-AC05-76RLO 1830. A portion of this research was performed using EMSL, a national scientific user facility sponsored by the DOE’s Office of Biological and Environmental Research and located at PNNL.

